# GenDiMug: a generic inclusive multigraph framework to analyze genomic organizations and -omics data at multiple scales

**DOI:** 10.64898/2026.10.06.756533

**Authors:** Jérôme Teulière, Philippe Lopez, Eduardo Corel, Briardo Llorente, Eric Bapteste

## Abstract

After decades of investigation, analyses of genome organization remain challenging. This is because genome descriptions are very information-rich and usually rendered as a linear map of genomic features corresponding to an ordered list of position intervals on a nucleic acid macromolecular backbone. This formalism, while accurate, is not fully optimized to mathematically analyze all genome structural properties, in particular the relations between genomic features, nor to integrate large amounts of -omics data, such as transcriptomes, epigenomes, proteomes, meltomes or chromatin accessibility data, which are increasingly available. We therefore developed GenDiMug, a tool that uses a network representation of the genome as a backbone, providing a natural framework to integrate spatial and ontogenetic relationships between genomic features and multi-omics data. We illustrate how this generic approach can be applied to all sorts of genomes, both to visualize genomic topology at contig and chromosomal scales at base-pair resolution, and to analyze genome positional effects using various graph metrics. Using GenDiMug, we further reveal clumping of genes with similar expression levels in eukaryotic genomes and their possible circadian regulation in a photosynthetic alga.

## Introduction

Multiple selective pressures have been shown to affect genomes’ organization (1–3). Therefore, systematic analyses with potential to unveil local to global structures in genomes can greatly enhance knowledge on genomics, developmental biology and evolutionary biology. Accordingly, affordable whole genome sequencing has led in the last 30 years to a wealth of genomic data that has been used as a research tool to, for instance, link the details of a genome sequence to medical outcomes or extract evolutionary insights along the Web of Life. For querying purposes, genome sequence and feature annotations are usually rendered as a linear map of genomic features, including e.g. Open Reading Frames (ORFs), intergenic regulatory regions binding to regulatory molecules, or DNA repeats, corresponding to an ordered list of position intervals on a nucleic acid macromolecular backbone. While this formalism has proven convenient, it is not optimized for mathematical analyses of genome structural properties. In living organisms, most biological processes require the interaction of spatially positioned features of genomes with external regulatory complexes (including complexes leading to chromatin compaction, replication, or epigenetic regulation of gene transcription), or three-dimensional rearrangements of the genomic macromolecule itself to bring distant features in close proximity, for which the map formalism complicates representation and modelization. In addition, beyond genomic data, large amounts of -omics data, such as transcriptomes, epigenomes, proteomes, meltomes or chromatin accessibility data (e.g. ATAC-seq) data, are becoming increasingly available, and easing their integration with genomics data is expected to enhance models describing how biological processes are influenced by the organization of genomes and how such processes affect in return genome structures in the course of evolution.

Precisely, integration of some of this biological complexity and data could be performed using a network representation of the genome as a backbone. However, whereas other large -omic datasets have been (largely independently) explored using network representation to derive systemic insights (e.g. transcriptomic data represented as Gene Co-expression Networks or Gene Regulatory Networks, proteomic and interactomic data as Protein-Protein Interaction networks, homolog genes as Sequence Similarity Networks), the structure of genomic data, which could provide a natural framework to integrate and structure multiple-omics data with genomics properties, has rarely been modelled as networks of relationships. Granted, attempts have been made to integrate human genomic (as well as proteomic) data as knowledge graphs to empower precision medicine, by making explicit the relationships between genomic features and medically relevant parameters recorded in various relational databases (4–10). However, topological patterns and genomic structure were not investigated in a generic and species-agnostic fashion using a network formalism, applicable to all genomes across the Web of Life, from those of mobile genetic elements to those of multicellular organisms, due to a current lack of powerful, inclusive tools.

To address this limitation, we developed GenDiMug, a tool able to generate genomic feature networks from genome sequences and annotations, allowing to bridge spatial and ontogenetic relationships between genomic features with other types of -omic data. This genomic features network representation provides a visualization tool for genomic topology at contig and chromosomal scales, allowing direct representation of dynamic changes affecting genomes over time with, when required, the precision level of one base pair. Moreover, it can also serve as an inclusive comparative framework to ease analyses of positional effects in genomes. As a proof of principle, we performed various analyses using graph metrics such as assortativity and shortest paths analyses to identify spatial structuration in gene functions and expression, and detected significant clustering of essential genes both in prokaryotic and eukaryotic genomes.

By mapping co-expression data onto edges of genomic networks, we also identified co-expressed genomic segments, including known and novel candidate operons, mobilized at specific stages during the archaeal cell cycle. We further detected domains on eukaryotic chromosomes composed of clusters of genes with similar expression levels, forming genomic regions under common evolutionary pressures (11–13) and regulations, whose patterns of expression were, in a photosynthetic species, strongly correlated with the progression of the circadian clock. Such iso-expressed chromosomal domains, in particular those with conditional timings of expression, differ from operons, but are compatible with other forms of optimized clumpings of independently expressed genes along specialized eukaryotic chromosomes that modulate gene expression levels via ‘loud’ and ‘quiet’ regions.

## Materials and Methods

### Genomic feature network building

We developed GenDiMug (https://github.com/TeamAIRE/GENDIMUG), a python script able to parse a genome annotation file (in the GFF3 format) and to build a genomic features network as a directed multigraph, i.e. a graph composed of multiple types of directed edges connecting multiple types of nodes (Fig. 1A). In such a graph, nodes are genomic features, and edges represent the ontogenetic (‘parent to child’) relationships between these features (Fig. 1B) and spatial (relative to their positions on the genomic sequence) (Fig. 1C).

**Figure 1.**
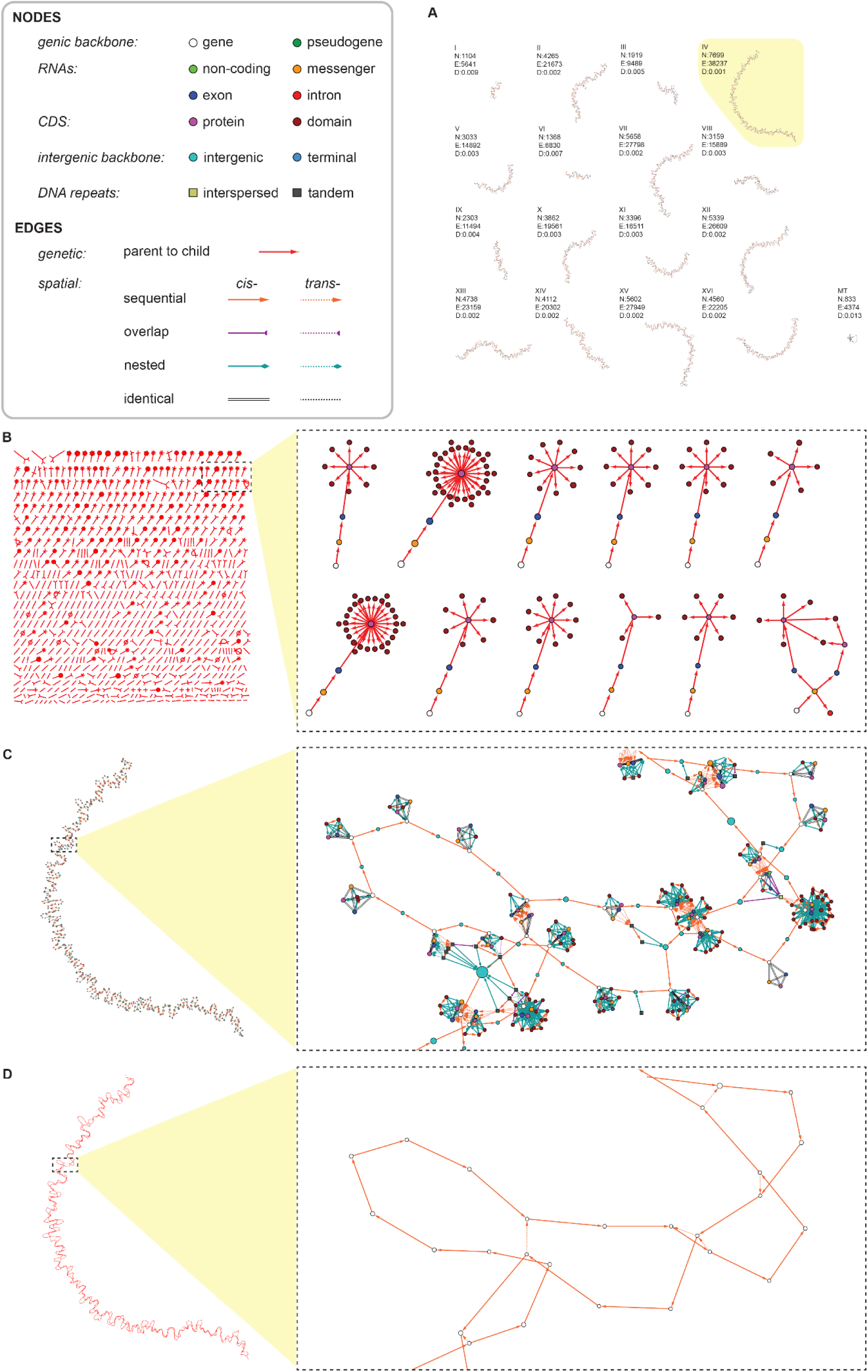
Genomic annotations can be converted to a genomic network. *S. cerevisiae* genome annotations (Refseq assembly GCF_000146045.2_R64) were used to build the corresponding genome feature network, in which nodes are genomic features (colored by feature types), and directed edges represent the ontogenetic (‘parent to child’) and spatial (relative to their positions on the genomic sequence) relationships between these features, in *cis* (on the same strand) or in *trans* (on opposite strands). (A) Whole genome network representation. Each chromosome (numbered in roman numerals, except for the mitochondrial genome indicated MT) corresponds to a connected component of feature nodes. The numbers of nodes (n) and edges (e) as well as the density (d) are indicated for each component. The chromosome IV is highlighted. (B-D) Details of the chromosome IV networks, segmented into (B) the genetic network, (C) the spatial network and (D) the simplified gene spatial network (GSN). (B-D) a magnification allowing to visualize the precise network structure is presented in the right panels, with the same genes in (C) and (D).

The minimal set of input files comprises a file containing the genomic annotation and a nucleic fasta file corresponding to the genomic sequence, which are readily provided for sequenced genomes recorded in the GenBank (https://www.ncbi.nlm.nih.gov/genbank/), RefSeq (https://www.ncbi.nlm.nih.gov/refseq/) or Ensembl (http://www.ensembl.org) databases. Additional features can be added to build a more complex network, such as DNA repeats, protein domains and motifs, or any custom genomic features annotated in the GFF3 format. Finally, the user can also provide custom annotations in the GFF3 format.

To build a graph, the script also infers features implicitly described by the genomic annotation (intergenic, terminal regions and introns) and defines each feature genomic positions as a list of position intervals, which allows to include fragmented features, i.e. features mapping to several position intervals on the genomic sequence (e.g. trans-spliced mRNA, or protein and proteic motifs encoded by multiple exons). It then generates annotation files for inferred and externally imported features that can be used to complement the reference genomic annotation.

Edges and nodes are associated to attributes. Edge attributes, contained in the network edge list, include the distances and overlap lengths (in bases) between features as well as the edge type (e.g. ‘cis-nested’, ‘reverse-complementary’…), and the edge class (e.g. ‘gene_cis-nested-gene’, ‘exon_cis-sequential_intron’, …) which allow to filter edge lists. Node attributes are exported into an annotation table comprising the original data from genomic annotations, the new list of positions and the GC percent for each feature.

First, the script collects the genomic positions of ORFs (feature: ‘gene’, including pseudogenes) for each listed contig and DNA strand (i.e. ‘+’ or ‘-’) to infer the complementary positions of intergenic regions between non-overlapping ORFs. It then creates linear backbone graphs for each strand of each contig by alternating ORF and intergenic features (or ORF and ORF when directly adjacent) linked by directed edges indicating sequential spatial relationships in cis (’cis-sequential’), in the order of their respective genomic position intervals. Terminal regions are inferred at the extremities of each of these strand graphs and from the contig positions. In the case of circular contigs (as indicated by the flag ‘is_circular=True’ in contig descriptions in the GFF annotation file format), terminal regions for each strand are unique nodes connecting the two ends of each strand graph. In addition, overlapping or nested ORFs are connected to the backbone by other spatial edges (’cis-overlap’, ‘cis-nested’). Intra-ORF features (RNAs, exons, CDS) are then connected to the ORF features in the network with similar cis-spatial edges. Exonic positions, when listed in the annotation file, are used to infer intronic positions and exons and introns are connected in sequence by cis-sequential edges. Finally, sequential, overlapping and nested relationships are computed in trans between features on the two strands. For efficiency, the script uses the interval tree data structure (https://pypi.org/project/intervaltree/) to identify overlapping and nested features, but custom code has been added to position features corresponding to a single position relative to other features. Features with different identifiers but with the same genomic positions are connected by undirected ‘cis-identical’ (in *cis*) or ‘reverse-complementary’ (in *trans*) edges. Additional features are connected to the genomic backbone by spatial cis- and trans-edges.

Intra-ORF features are also connected by genetic directed edges from ‘parent’ to ‘child’ (as indicated by the GFF flag ‘Parent’), with the following modifications relative to the relationships described by the genome annotation: Exons, instead of mRNAs, become the parents of the CDS parts; trans-spliced ORFs are explicitly fragmented as gene parts and are all parent to the respective trans-spliced mRNAs; if annotated by an external software such as Interproscan in the GFF3 format, protein domains and motifs become the ‘children’ of the CDS to which they belong. Each ORF has thus an associated parental subgraph oriented from gene (or gene part) to RNA, RNA to exons and to introns, exon to CDS (or in the case of prokaryotic annotation, directly from gene to CDS) and CDS to protein domain/motif (Fig. 1B). To limit complexity, nodes for identical features of the same type are collapsed into a single node and nested DNA repeats are removed. In total, the complete networks can be composed of 12 node types: gene; pseudogene; rna; exon; intron; cds; domain; intergenic; terminal; tandem repeat; interspersed repeat; additional feature; and 9 edge types: ontogenetic; cis-sequential; cis-overlap; cis-nested; cis-identical; trans-sequential, trans-overlap, trans-nested, reverse-complementary. Four network simplification options are implemented (Spatial network without children features; gene-only spatial network backbone with additional features; spatial gene-only network, ontogenetic-only network).

### Genomic and transcriptomic datasets

Complete genomic sequences (nucleic and protein fasta files) and annotations (GFF3 file) were downloaded from the RefSeq (https://www.ncbi.nih.org/refseq/) or GenBank (https://www.ncbi.nih.org/genbank/) sequence databases to generate genome feature networks for the species and accession codes listed in Supplementary Table S1. Additional features were added to complement the base genomic annotation files using the following software: protein domains and motifs were annotated in the GFF3 format using standalone interproscan v5.60-92.0 (40) with applications Pfam, Phobius and PRINTS; tandem repeats were detected using the pipeline Tandem Repeats Finder v4.09 (41) using parameters Match = 2, Mismatch = 7, Delta = 7, PM = 80, PI = 10, Minscore = 50, MaxPeriod = 2000, and Tandem Repeats Analysis Program v1.1 (42) using default parameters with -id 70; interspersed repeats were annotated in the Stockholm format using RepeatModeler v2.0.4 (43) using default parameters and the - LTRStruct flag.

The following transcriptomic datasets were downloaded from the Gene Expression Omnibus repository (https://www.ncbi.nlm.nih.gov/gds): *C. rheinardtii* GSE71469 (44) and GSE112394 (45), *S. cerevisiae* GSE90008 (unpublished), GSE240513 (46) and GSE168699 (47). *S. islandicus* transcriptomic dataset was described in (15). Raw counts were normalized to TPM (Transcripts Per Kilobase Million) or RPKM (Reads Per Kilobase Million). Normalized counts were further classified by quantiles based on the values in the whole dataset before network analysis.

### Categoric annotations

*S. islandicus* gene essentiality annotations were mapped from the DEG database (48). *E. coli* K12 essential genes were those described in (49). *S. cerevisiae*, *S. pombe,* and *C. elegans* essentiality annotations were those from the OGEE v3 database (50). To account for the possibility that deletions targeting more than one gene (and thus multiple adjacent genes) in *C. elegans* may lead to a lethal phenotype, the allele description file was retrieved from Wormbase (https://wormbase.org, release WS295) and, as a correction, the essentiality annotation was removed for genes whose loss-of-function mutant lethal phenotype was only associated with multigenic deletion mutations.

Cell cycle genes from these species and from *D. melanogaster*, *H. sapiens* and *M. musculus* were defined using AmiGO 2 (https://amigo.geneontology.org/amigo) by their association with the Gene Ontology term GO:0007049 ‘cell cycle’. *S. islandicus* ArCOG annotations were derived from the ArCOG database (16) and *E. coli* COG annotations were derived from the COG database (51). *S. islandicus* gene conservation was described in (15). Aging-associated genes were those recorded in the GenAge database (https://genomics.senescence.info/genes/index.html) (52).

### Label correction for described gene clustering

To correct for genomic proximity either due to gene evolution by duplication or transcript polycistronic organization, categoric node labels were removed from genes connected through a shortest path if they were expected to cluster together, either as arrays of ancestrally-related genes or as described components of polycistronic mRNAs and operons. To determine which genes may belong to clusters of ancestrally-related/homologous genes, a diamond blastp all-against-all search was conducted with the proteins encoded by the genes from the subset of labelled nodes in order to define gene families by sequence similarity network filtering (minimum 30% identity and 80% mutual coverage), using the scripts cleanblast and familydetector (53). The *C. elegans* gene-operon mapping was extracted from the genomic annotation file c_elegans.PRJNA13758.WS295.annotations.gff3 downloaded from Wormbase (https://wormbase.org, release WS295), and the list of known *D. melanogaster* genes belonging to polycistronic mRNAs was retrieved from Flybase (https://flybase.org, release FB2025_01) using the gene-associated Sequence Ontology term SO:0000690 ‘gene_with_polycistronic_transcript’ as query for a Vocabularies search. *S. cerevisiae* genes occasionally expressed as bicistronic transcripts were described in (54).

### Assortativity and shortest paths analysis

After conversion of the directed gene-only multigraph to an undirected graph, the functions attribute_assortativity_coefficient and numeric_assortativity_coefficient from the networkx python package (https://networkx.org/) were used to compute assortativity coefficients for node categoric and numeric labels, respectively (14). For transcriptomic data, numeric node labels were the quantile associated to the normalized read count value for the distribution of all values in a dataset; similar assortativity patterns were observed with 10 (deciles), 50, 100 (centiles) or 1000 quantiles (data not shown). In practice, centiles were used to produce assortativity heatmaps, and deciles for the calculation of edge proportions (Supplementary Fig. S7) and for the visualization of labelled gene clusters (Supplementary Movie S1, Supplementary Fig. S6), in order to reduce the number of labels. The shortest path length between each pair of genes on each contig was determined using function all_pairs_shortest_path_length from the networkx package, with a shortest path length cutoff set at 50 edges. Specific analysis of these lists of shortest paths was performed with custom python scripts to calculate the numbers of direct edges (paths of length 1) between labelled nodes of interest and the average shortest path length for each labelled node to the closest similarly labelled node.

### Intersection of co-expression and spatial edges

Network intersection of *S. islandicus* genomic features networks with gene co-expression networks was performed by computing the set intersection between (1) the set of gene pairs connected by edges corresponding to significant consensual positive correlations in each gene co-expression network associated to a cell cycle phase (15) and (2) the set of node pairs connected by a spatial edge from the gene spatial network.

### Genomic network visualization

Networks were visualized using Cytoscape (https://cytoscape.org/). To assemble animations of these networks for time-series of transcriptomic data, images of the same regions of the networks were collected for each time-dependent sets of annotations, and ImageJ (https://imagej.net/software/imagej/) was used to animate and label the stack of network images.

### Statistical tests

Empirical p-values were determined by label permutation tests as described in (55) and adjusted using the Bonferroni procedure. For categoric labels, shuffling of labels was performed per contig graph to account for the non-homogeneous distribution of labelled nodes on each chromosome. For numeric labels, nodes labels were shuffled per whole genome graph. To determine whether gene expression assortativity coefficients (r) significantly changed in transcriptomic time-course datasets, time series were split in 2 or 3 consecutive series and the corresponding distributions of r were compared, by contig, using a Wilcoxon U-test, for which p-values were adjusted using the Bonferroni procedure.

## Results

Annotations of sequenced genomes describe both spatial relationships between genomic features on the physical genomic macromolecule (DNA or RNA), commonly rendered as a genomic map in a genome browser, and ontogenetic (i.e. ‘parent to child’) relationships between each Open Reading Frame (ORF), its transcripts and eventual translated products. To exploit these underlying systemic relational structures, these data can also be represented as a graph, which could be simply used as a knowledge graph, or be analyzed using graph tools and metrics in order to derive novel biological insights. Furthermore, a genomic feature network can be used as a generic, species-agnostic and expandable network backbone to integrate multiple - omics datasets as new nodes, edges, node or edge attributes or network topology modifiers, e.g. to identify patterns in the topology of the graph or changes in graph metrics between various conditions or across species. So far, to the best of our knowledge, such generic representation of genomes as networks has not been implemented, nor exploited.

We developed GenDiMuG, a python script with the aim to convert <u>Gen</u>omic annotations and sequences to a <u>Di</u>rected <u>Mu</u>lti<u>G</u>raph. This graph comprises multiple types of nodes (genomic features) connected by multiple types of directed and undirected edges (corresponding to spatial and ontogenetic edge types), and its connected components correspond to contigs, or chromosomes in complete genomes (see M&M, Fig. 1). To test this tool on annotated genomic data of various sizes, we selected 23 datasets from viral (DNA viruses), prokaryotic (eubacteria and archaea) and eukaryotic (protists, plants, fungi and animals) genomes, computed additional annotations (protein domains and DNA repeats) and produced the corresponding genomic networks, which sizes ranged from ∼50 up to ∼20M edges depending on the number of annotated features and the richness of this annotation (Supplementary Table S1). By construction, due to their overall linear structure, genomic feature networks are edge-sparse with a number of edges corresponding to roughly 3 to 7 times the number of nodes and thus have a low density (ratio of observed edges to the number of possible edges for a given network) (Supplementary Table S1). These networks can be segmented as purely ontogenetic or spatial edges (Fig. 1A-C) and also be simplified to perform specific analyses (Fig. 1D). To illustrate the usefulness of genomic networks to perform genomic neighborhood analysis, we used the simplified gene spatial networks (GSNs) derived from genomic annotation of the archaeal *Saccharolobus islandicus* and bacterial *Escherichia coli* prokaryotic genomes, and of the eukaryotic genomes from model species *Caenorhabditis elegans*, *Drosophila melanogaster, Mus musculus*, *Homo sapiens, Chlamydomonas reinhardtii* and *Saccharomyces cerevisiae* in the examples showcased below.

### Assortative mixing and shortest-paths analyses with categoric node labels

In addition to feature attributes found in the genomic annotations (e.g. intron, exon, gene, …), the genomic feature nodes in genomic networks can be associated to various attributes, either categoric (i.e. a text label describing a property of the feature, e.g. a phylogenetic origin) or numeric (i.e. a numeric value associated to a feature, be it a rank or a measured value for a given metrics, e.g. an expression level). In such a labelled network, the assortative mixing of nodes, i.e. the similarity of neighboring nodes in the graph with respect to a given attribute, can be measured using the assortativity coefficient *r* (Supplementary Fig. S1). This assortativity coefficient can either take negative values comprised between −1 and 0 (for a disassortative network in which similarly labelled nodes tend to never be directly connected) or positive values between 0 and 1 (for an assortative network in which similarly labelled nodes tend to be increasingly directly connected) (14). Furthermore, by using label permutation tests, empirical p-values can be computed to test whether a measured value of *r* for a genomic network is significantly different than values obtained with randomized networks.

Assortativity analyses are typically relevant to uncover broad-scale patterns of the genomic organization, e.g. when features with similar origins or expression levels tend to clump together, forming detectable domains at specific chromosomal positions. To illustrate this, we first tested whether different categoric node labels showed an assortative mixing pattern in the GSNs from prokaryotic genomes. For the archaeon *S. islandicus*, labels indicating levels of gene evolutionary conservation in the Sulfolobales phylum (15) and ArCOG functional categories (16) were associated with moderate but significantly positive assortativity (r = 0.14 and r = 0.17, respectively; p < 0.05, label permutation tests) suggesting that conserved and functionally related genes tend to cluster in this phylum. Moreover, the gene essentiality-based assortativity coefficient was 0.35, indicating a strong tendency of essential genes to be direct neighbors in the graph. Similarly in the bacteria *E. coli* gene network, the essential genes and the COG categories distributions were also assortative, with significantly high assortativity coefficients of 0.3 and 0.16, respectively.

Such a structuration by gene categories is not surprising in prokaryotic genomes, known to undergo selective pressures (17–19) to optimize gene neighborhoods by clustering genes of shared functionality, oftentimes being co-transcribed into polycistronic mRNAs from operons in Bacteria and Archaea, as well as being concentrated in large-scale chromosomal compartments of highly and lowly expressed genes in Archaea (20, 21). Indeed, 374 of the 429 (87.2 %) *S. islandicus* essential genes are located in the highly expressed A genomic compartment.

Essential genes may be subject to evolutionary pressures maintaining a clustered organization of genes in the genome by functional similarity. This can be directly tested by extracting from the genomic network all the nodes labeled as essential and forming components of at least two nodes (one edge) from the network, and by estimating their functional similarity using ArCOG categories (16), showing the versatility of this multidirected network framework. In support of this hypothesis, for all pairs of nodes connected by a spatial edge in the subgraph formed by essential genes, 39.8 % of gene pairs were labelled with the same functional category (n = 251 edges) which is significant compared to proportions observed when the essential genes are randomly relabeled by shuffling their ArCOG category labels (p < 0.05, label permutation test). For *E. coli*, the assortativity coefficient value is consistent with the fact that 243 of 358 essential genes (67.9 %) were located in known operons, with 295 that were in the same operon as at least one other essential gene (82.4%), and 81 of the 150 pairs of essential genes connected by an edge in the network (54%) shared the same functional category, again a value significantly higher than with randomly shuffled COG category labels (p < 0.05, label permutation test).

By contrast, in eukaryotes, the annotated GSNs from *S. pombe*, *S. cerevisiae* and *C. elegans* were generally not significantly assortative for essentiality, indicating much less functionally structured gene neighborhoods, except for *C. elegans* chromosome I with a low but significant positive assortativity (r = 0.04, p = 0.006). However, locally assortative structures in the graph could also result from gene tandem duplications (implying the duplicated paralogs would remain essential) or association in polycistronic expression clusters such as operons (22).

Beyond assortativity analyses, many other network metrics can be used to investigate genomic structure. For instance, shortest paths, i.e., the minimum number of edges to connect pairs of labelled nodes (Supplementary Fig. S1C), will provide metrics of topological feature proximity in the network (Supplementary Fig. S2), rather than a physical mapping distance. Interestingly, the average shortest path lengths between essential genes or between genes with the same function were significantly lower than in label-shuffled graphs, even after correction for known causes (gene duplication or polycistronic transcripts, see Methods) of gene clustering (Supplementary Figs. S3, S4). This approach also detected significantly high numbers of direct edges between essential genes in graphs from *S. cerevisiae* chromosome XV, all three *S. pombe* chromosomes, and again, *C. elegans* chromosome I (Fig.2A-C, top plot panels). However, after correcting for known causes of gene clustering in these species, only *S. pombe* chromosomes displayed significantly high numbers of direct edges between essential genes (Fig. 2A-C, bottom plot panels). In addition, the average shortest path length between two essential genes was lower than expected by chance, even after correction, for *S. cerevisiae* chromosomes IV, V, VIII, XIV and XVI, and for all three *S. pombe* chromosomes. In *C. elegans*, significant proximity of essential genes in the graphs was seemingly solely due to colocalization in operons and clusters of ancestrally related genes (Fig. 2D-F). These results suggest that eukaryotes could, like prokaryotes, be under evolutionary pressures to maintain a loosely clustered organization of their essential genes (23).

**Figure 2.**
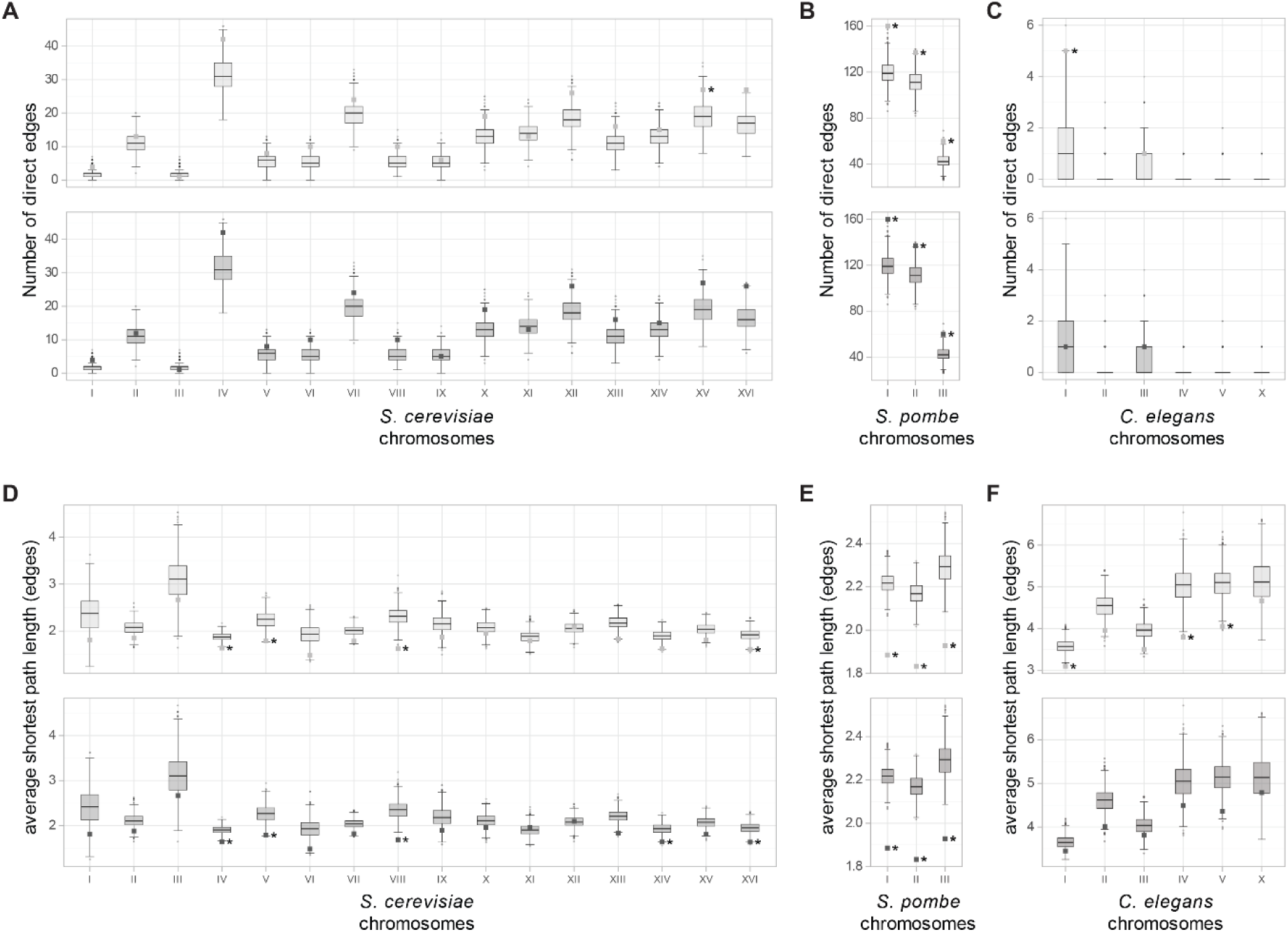
Shortest paths analysis in eukaryotic gene-only spatial networks labelled for gene essentiality. (A, B, C) Number of direct edges (shortest paths of length 1) and (C, D, E) average shortest path length between essential genes in (A, D) *S. cerevisiae*, (B, E) *S. pombe* and (C, F) *C. elegans* before (light grey) and after (dark grey) correction for known sources of gene clustering. The reference values are indicated by squares, and box plots represent the distributions of random values in label permutation tests. A star indicates when the reference value is significantly higher (A-C) or lower (D-F) than random values (p < 0.05 after Bonferroni correction, label permutation test).

Other categorical labels also showed unexpected limited but significant spatial structures in eukaryotic genomic graphs, even after corrections. For example, aging-associated genes were closer than expected by chance in *C. elegans* chromosome IV (12 direct edges, and a significantly lower average shortest path length between those genes) and *D. melanogaste*r chromosome 3L (significantly lower average shortest path length of ∼2.8Likewise, cell cycle-associated genes were clustered in *D. melanogaster* chromosome 2L (15 direct edges) and *M. musculus* chromosome 2 (8 edges).

### Connected components with similarly labeled edges

Attributes can not only be mapped on nodes of a genomic network but also on its edges. Such mapping can be important to determine if spatially close nodes contribute to common biological processes, or for instance are subjected to common regulations. To illustrate this, we labeled edges of genomic network of the archaeon *S. islandicus* depending on whether the two connected nodes had been reported to be co-expressed at a specific stage of its cell cycle in (15). Using the genomic network as a backbone immediately unveils sets of co-expressed neighbor genes, as this defines connected components (subnetworks connected by a co-expression edge but disconnected from the rest of the genomic network) for every cell cycle phase (Supplementary Fig. S5). Logically, such sets notably contained genes from all known *S. islandicus* operons except the archaellum. Interestingly, hundreds of additional components characterized by colocation and correlations in expression values were also identified by this method, potentially pointing to yet uncharacterized operons in this archaeon. Similar analyses based on edge mapping could be performed to analyze other properties, such as protein-protein interactions, by intersecting instead data from genomic networks with data from other types of networks, here protein-protein interaction networks.

### Assortative mixing of numeric node labels

Genes with related functions are not the only ones that may clump together in genomes. Genes with correlated transcription may also co-localize in pairs along chromosomes, as has been observed in several eukaryotic model species (24, 25). Patterns of co-expression as well as iso-expression (genes expressed at similar levels) of neighbor genes can be readily analyzed in genomic networks with numeric labels. To illustrate the use of genomic features networks to detect assortative patterns of genomic features associated to numeric labels, we mapped transcriptomic data on the nodes of the GSNs from two model eukaryotic organisms *S. cerevisiae* and *C. reinhardtii*. For these taxa, we selected bulk transcriptomic datasets obtained from homogeneous, cell-cycle synchronized populations of cells, suitable to detect time-dependent patterns of gene expression along genomes from time series generated to minimize inter-individual transcriptional variability while averaging the discrete cell-specific signals linked to transcriptional bursting that has been observed in single-cell datasets (26–28). We classified the normalized read counts associated with each gene in quantiles to normalize these values across samples and time points (see M&M). We used these quantiles as numeric attributes of the nodes of this yeast and this alga genomic networks, which provided a simple visualization of node labels in the networks.

Strikingly for both species and in each transcriptomic dataset tested, we observed significantly positive assortativity coefficients (with values between 0.1 and 0.3) for *S. cerevisiae* chromosomes VIII-X, XII and XIV (Fig. 3A) and for all *C. reinhardtii* chromosomes except 4, 11 and 14 (Fig. 3B). Therefore, neighboring genes on these chromosomes tend to display similar levels of expression in the corresponding genomic graphs. This exciting structure indicates that, convergently, some eukaryotic chromosomes harbor ‘loud’ and ‘quiet’ regions in terms of gene expression. This observation suggests that eukaryotes may spatially structure some of their chromosomes by expression level, which would suggest that gene positions may be optimized in genomes to cluster similarly expressed genes, without generally forming *bona fide* operons allowing the co-expression of functionally related genes as observed in prokaryotes, although polycistronic expression has also been described in eukaryotes (29–31), including in *C. reinhardtii* (32). Since the value of the assortativity coefficient *r* depends on neighbors being close in numeric values, rather than identical, it is likely that a positive *r* value reflects a continuum of expression levels across the graph. Indeed, groups of nodes with close levels of expression were consistently observed in animated visualizations of the assortative labelled graph from chromosome 6, compared to the weakly assortative graph from chromosome 14 (Supplementary Movie S1) and clusters of iso-expressed genes can be observed in segmented views of the corresponding connected components for *C. reinhardtii* (Supplementary Fig. S6).

**Figure 3.**
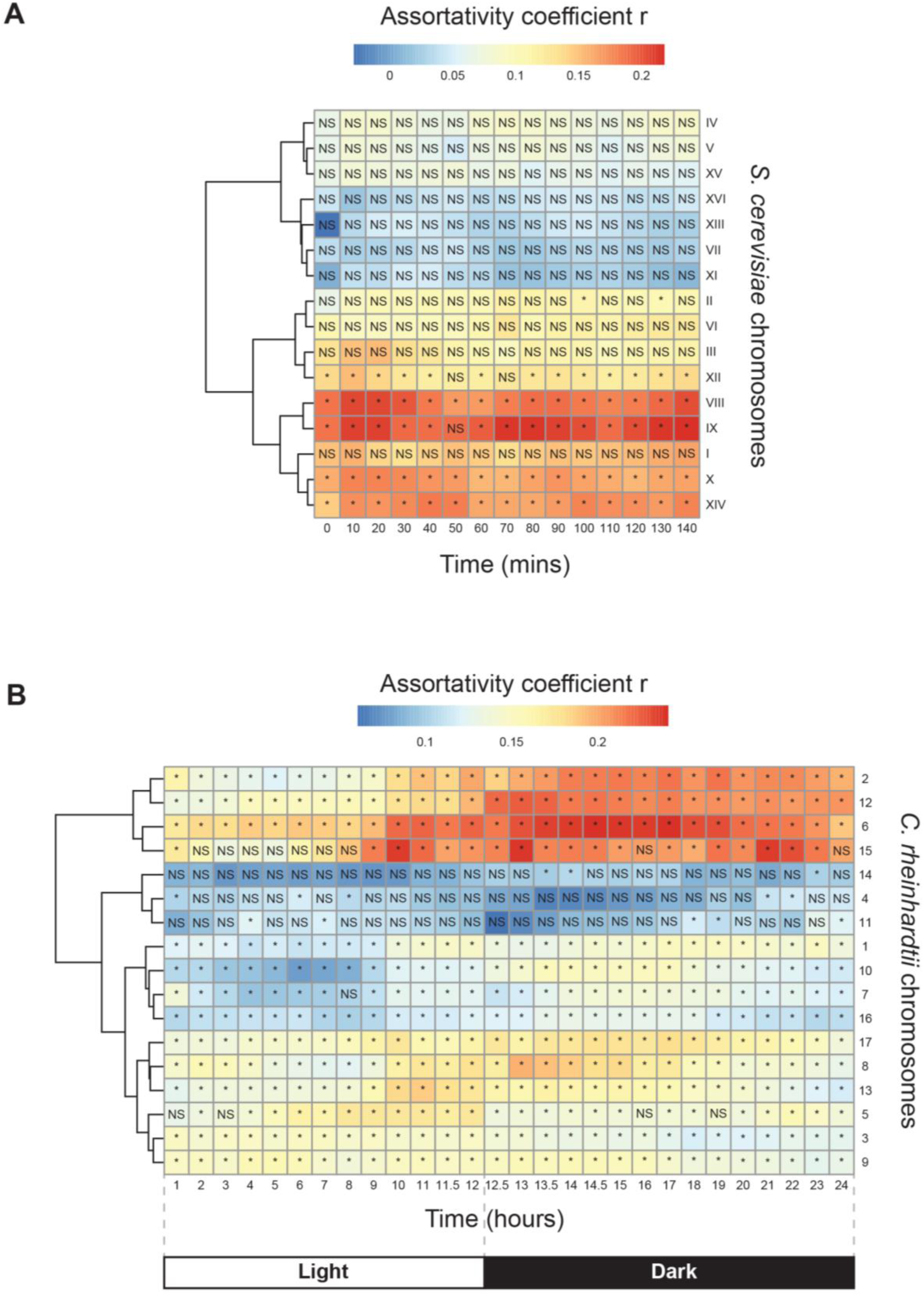
Assortativity analysis of gene-only spatial networks labelled for gene expression. (A, B) Heatmaps displaying the values of the assortativity coefficient *r* computed for numeric node labels in gene networks (the distribution of normalized read counts over the whole dataset defined centiles of expression, used as numeric node labels) for each chromosome and each time point from transcriptomic datasets produced by sampling synchronized cultured cell populations of (A) the yeast *S. cerevisiae*, every ten minutes over one cell cycle (47) and (B) the unicellular green alga *C. reinhardtii*, every hour or half-hour over a circadian cycle with a 12 h photoperiod, indicated with the boxes Light and Dark (44). A star indicates a significantly high value of r (node label permutation tests, p-value correction by the Bonferroni procedure). NS: not significant. Chromosomes were clustered by similarity of their temporal assortativity for gene expression. Chromosome names in bold typeface indicate chromosomes structured by gene expression (in black), and by gene expression and time (in red). *S. cerevisiae* mitochondrial and *C. reinhardtii* mitochondrial and plastidial chromosomes are missing because their genes were not available in the published datasets.

In addition, in the *C. reinhardtii* dataset, this assortative structuration of gene expression is changing over time (in particular for Chr. 2, 6, 12 and 15), likely in association with circadian rhythms and light-induced photosynthesis (Fig. 3B). Thus, ‘loud’ and ‘quiet’ genomic regions are not active at the same time of the day. For the *C. reinhardtii* dataset, we calculated for each chromosome graph the proportions, relative to the total number of edges in each graph, of iso-expressed edges linking two nodes from the same expression decile (Supplementary Fig. S7). Strikingly, the sum of the proportions of such edges was stable over time (between ∼12 and 20% of all edges), but most chromosomes displayed a decrease in the proportion of edges connecting nodes with low or no expression (decile 1) under light, and an increase of this proportion in the last hours of the dark period, presumably again reflecting circadian and light-induced expression patterns. Why this is the case is beyond the scope of our paper, but one may imagine some coupling between gene expression intensity and the metabolic activity of this alga in presence or absence of light, as highly iso-expressed genes are active during the day, whereas lowly iso-expressed genes are active at night, and that, as previously shown in other eukaryotes, some common spatial regulation (33), possibly a restriction or an ability to access these regions of the chromatin, contributes to the silencing or to the activation of these clusters of genes (34).

Similar chromosome-scale assortative gene expression was also observed using two more *S. cerevisiae* and one more *C. reinardtii* transcriptomic datasets in time series, obtained with synchronized population (Supplementary Fig S8), confirming that the above results are not an artefact, but reflect a robust, convergent property of eukaryotic genomes.

## Discussion

Genomes are usually represented using linear maps of genomic features, but this formalism supports less explicit mathematical treatments of the genomic organizational features than the derived multi-digraph formalism that we introduced here and that we called genome features networks. There are indeed several advantages to rely upon networks to investigate genomic data. First, networks provide a natural, generic powerful framework to visualize these ontology-rich data, using diverse types of nodes and of edges. Moreover, a complex genome feature network can always be decomposed (by type of edges) or simplified (by type of nodes) for specific analysis (e.g. gene-only spatial network allowing the mapping of transcriptomic data to nodes), or combined with other types of networks (e.g. protein-protein interaction network, sequence-similarity network, gene co-expression network, etc.) by attributing labels to the edges of genome feature networks. Incidentally, these networks can be used as a curation tool of genomic maps, as they highlight gaps or discrepancies in annotations (e.g. childless genes i.e. genes with no annotated transcript or CDS, inconsistent CDS positions and protein sequences in genome protein fasta file), or inconsistent ontogenetic flow (e.g. gene to exon direct link).

Second, this network formalism allows for quantitative and statistical analyses of the structures within genomes, and even (although this was not done here) to compare these structures across genomes. We illustrated the inclusiveness of this framework using it to address a diversity of research questions as applications, such as the clumping of essential or aging-associated genes along chromosomes, operon detection and prediction, or the identification of iso-expressed domains on some chromosomes, sometimes under possible circadian control, from transcriptomic time-series mapped onto genome feature networks. Gene clustering by functional similarity is characteristic of prokaryotic genome organization, and thought to be exceptional in eukaryotic genomes (25, 35–37). Indeed, we largely observed assortative mixing of nodes by function in *E. coli* and *S. islandicus* genomes, but also observed that genes whose null phenotype in experimental settings leads to lethality were clustered as well, both in prokaryotic and some of the model eukaryotic genomes we analyzed. Such an unexpected observation may be explained by phenotypic annotation strategies that attribute a lethal phenotype linked to a large deletion to all the genic elements it encompasses. Although we corrected for these possible artifacts in *C. elegans* phenotypic annotations (see Methods), and we checked that *S. cerevisiae* and *S. pombe* lethality data originated from single-loci mutations and precise single-ORF targeted deletions genes (38, 39), it is possible that biases in annotations remain and could explain the observed phenotypic similarity of neighboring genes. Alternatively, an evolutionary pressure to cluster essential genes could make sense if gene expression of neighbors tends to be similar. Finally, spatial clustering of genes associated with diseases or aging may point towards critical genomic regions with potential for medical applications.

Although previous transcriptomic studies in eukaryotic model species have shown that adjacent pairs/triplets of genes are co-expressed (reviewed in Michalak 2008), co-expression was defined by creating matrices of Pearson correlations for all gene pairs over whole datasets of samples and conditions, thus identifying correlations that are always observed regardless of experimental conditions, and of levels of expression. In contrast, our approach shows that genes of similar expression levels, as indicated by their expression quantile numeric node labels in gene-only networks, are often, on a chromosome basis, not randomly positioned in the genome but rather direct neighbors, and that this clustering is dynamic in time-course experiments. Except expression quantiles computation, this similarity was not defined over all samples of a dataset but observed independently in each sample by mapping normalized read counts on the nodes of a reference gene network. To our knowledge, this has not been observed by previous studies and represents a benefit of the network-based approach to analyze gene expression. To sum up, genome features network can provide a kind of generalizable spatial network to observe the dynamics of co-expression clusters, while using network tools (assortativity, short paths analyses, network filtering by quantiles of expression) and directly visualizing gene expression in a genomic context, potentially in relation to other genomic features.

Still, beyond transcriptomes, many other mappings would be of course possible, and epigenomics data, ATAC-Seq, protein-protein interactions and labels reflecting the phylogenetic origins of the components of the genomes (e.g. nuclear genes from an endosymbiont in the case of eukaryotes or laterally acquired genes in the case of prokaryotes) could all be integrated with genome features networks. Amongst these future promising directions, analyses of the epigenetic and transcriptomic dynamics along the genome could inform studies from developmental biologists; and comparative studies of genomes from closely related species (e.g. non-human apes and humans), or of genomes from bacteria undergoing experimental evolution or of synthetically scrambled genomes generated by synthetic biologists using genome features networks may illuminate important aspects of genome organization and regulations and its conservation across species.

Supplementary Data are available at NAR online.

## Acknowledgements

We thank Miguel V. Gomez-Raya-Vilanova, Virginija Cvirkaite-Krupovic and Mart Krupovic for helpful discussions on archaeal genetics. BL gratefully acknowledges support from the ARC Centre of Excellence in Synthetic Biology, Macquarie University, Bioplatforms Australia, Twist Bioscience, the National Health and Medical Research Council (NHMRC), the Ramón Areces Foundation, and the Allen Foundation.

## Author contributions

Jérôme Teulière (Conceptualization, Data curation, Methodology, Software, Validation, Visualization, Investigation, Formal analysis, Writing – original draft, Writing – review & editing), Philippe Lopez (Conceptualization, Writing – review & editing, Funding acquisition), Eduardo Corel (Methodology, Software), Briardo Llorente (Conceptualization, Funding acquisition, Writing – review & editing, Project administration) and Eric Bapteste (Conceptualization, Funding acquisition, Formal analysis, Writing – original draft, Writing – review & editing, Project administration).

## Data availability

The data underlying this article are available in the article and in its online supplementary material.

