## Supplementary Figure 1 for "GenDiMug: a generic inclusive multigraph framework to analyze genomic organizations and -omics data at multiple scales"

**A**

$$r = \frac{\Sigma_i e_{ii} - \Sigma_i a_i b_i}{1 - \Sigma_i a_i b_i}$$

**B**

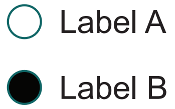


Disassortative network (r = -0.58)

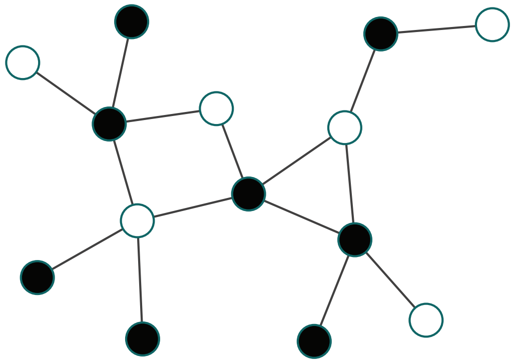

Assortative network (r = 0.28)

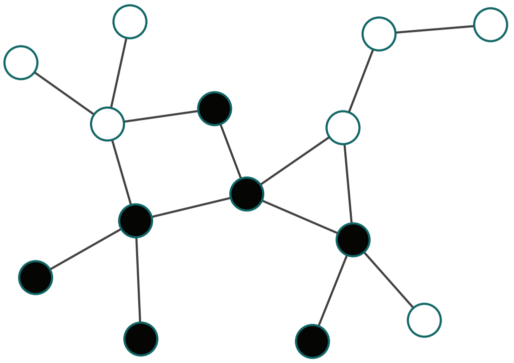

**C**

Shortest path between two similarly labelled bodes (length = 3 edges)

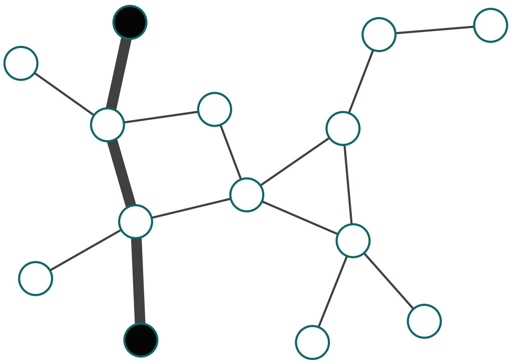
