## Supplementary figures and images for "GenDiMug: a generic inclusive multigraph framework to analyze genomic organizations and -omics data at multiple scales"

### Supplementary Figure 2

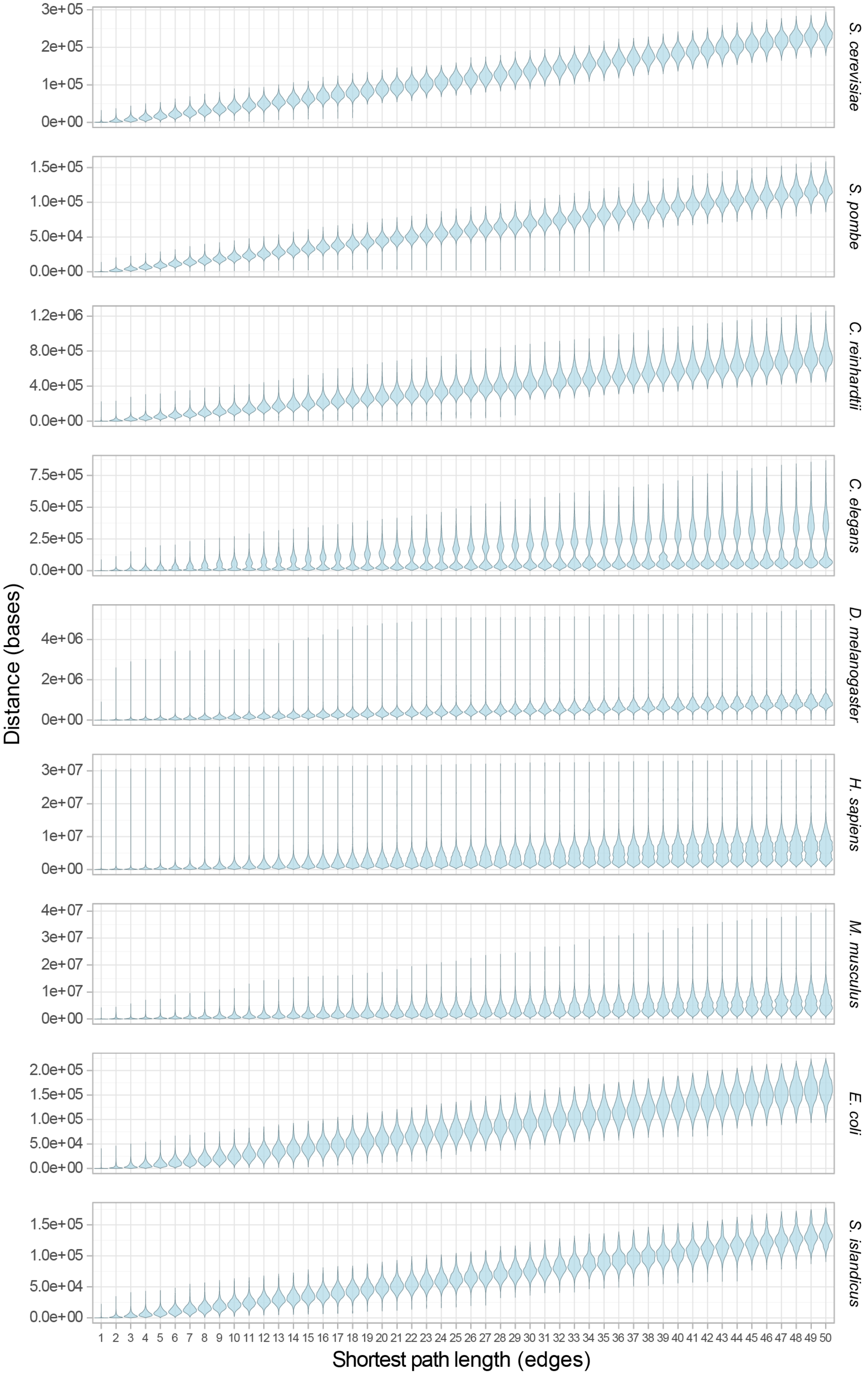

### Supplementary Figure 3

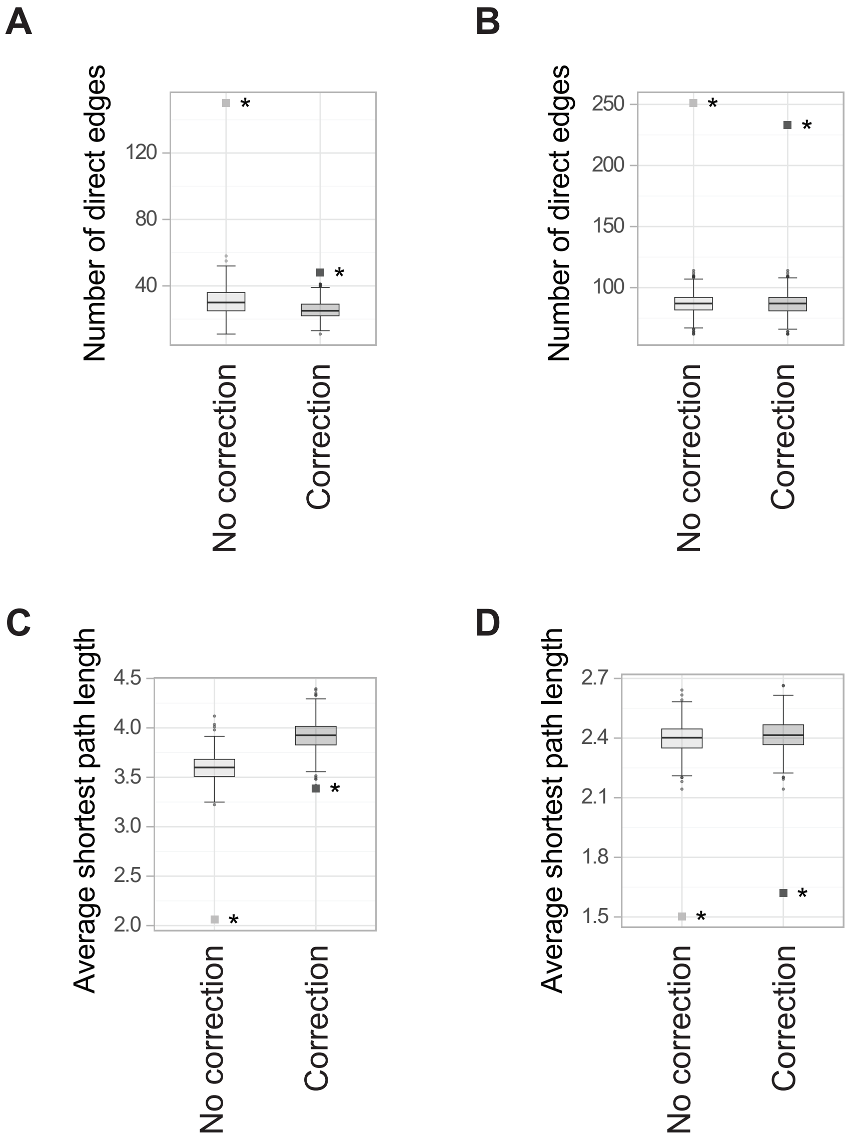

### Supplementary Figure 4

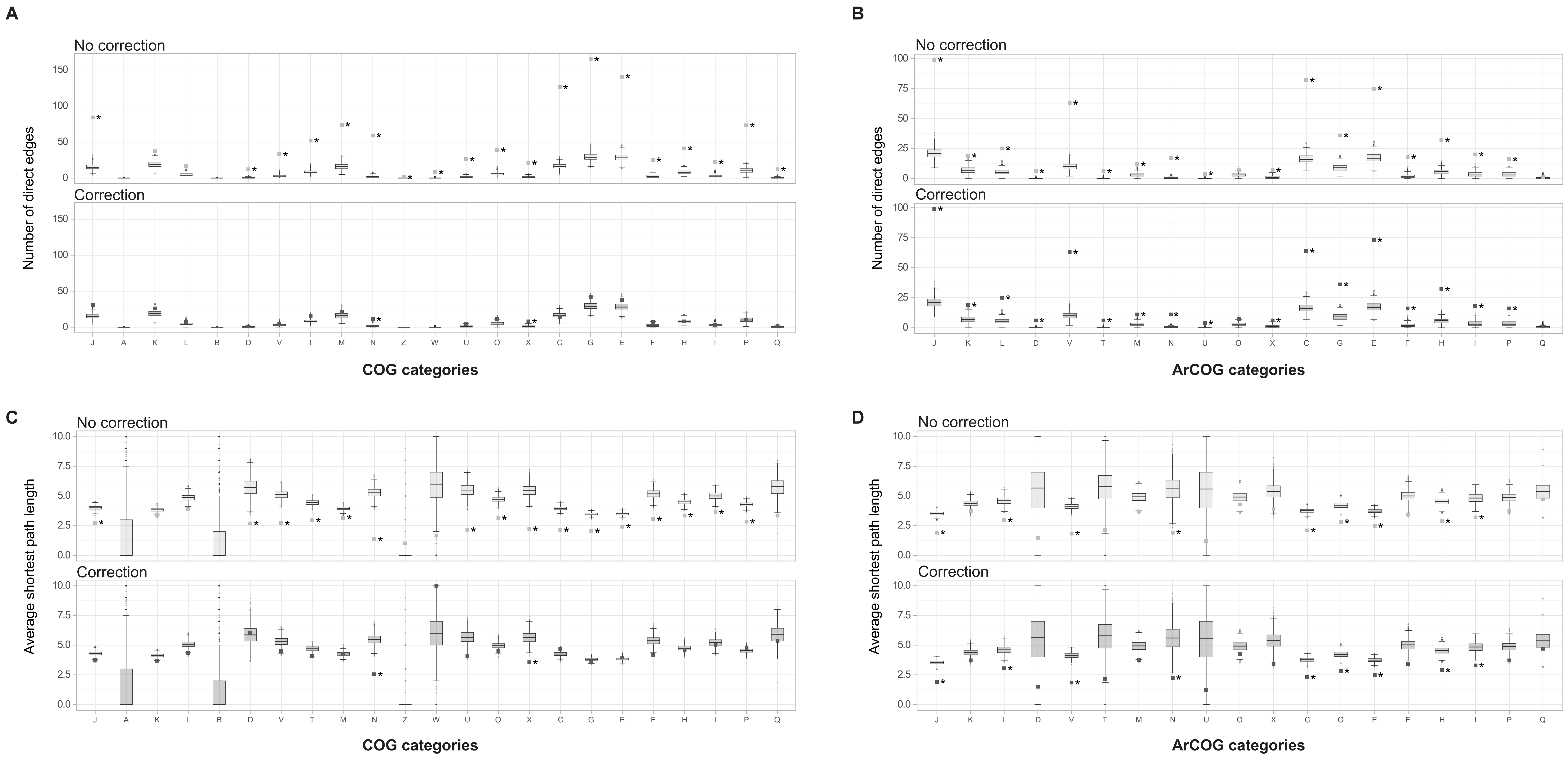

### Supplementary Figure 5

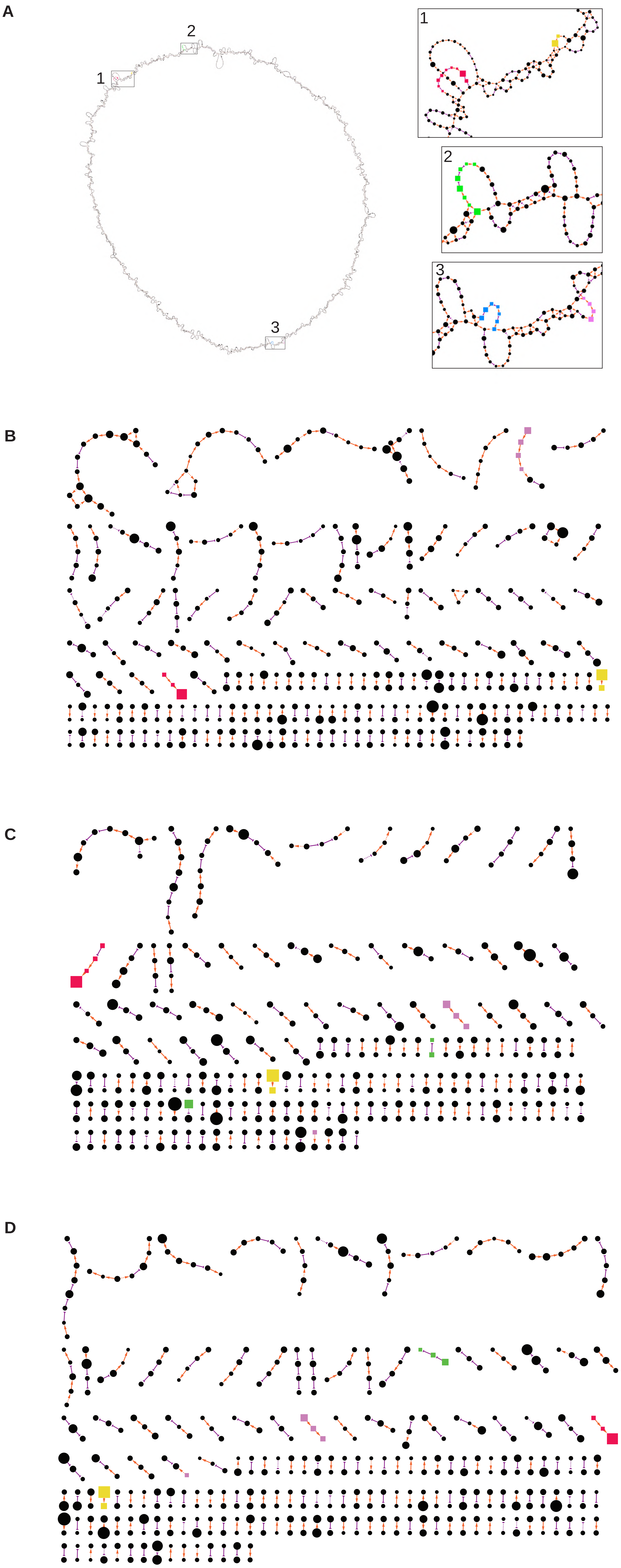

### Supplementary Figure 6

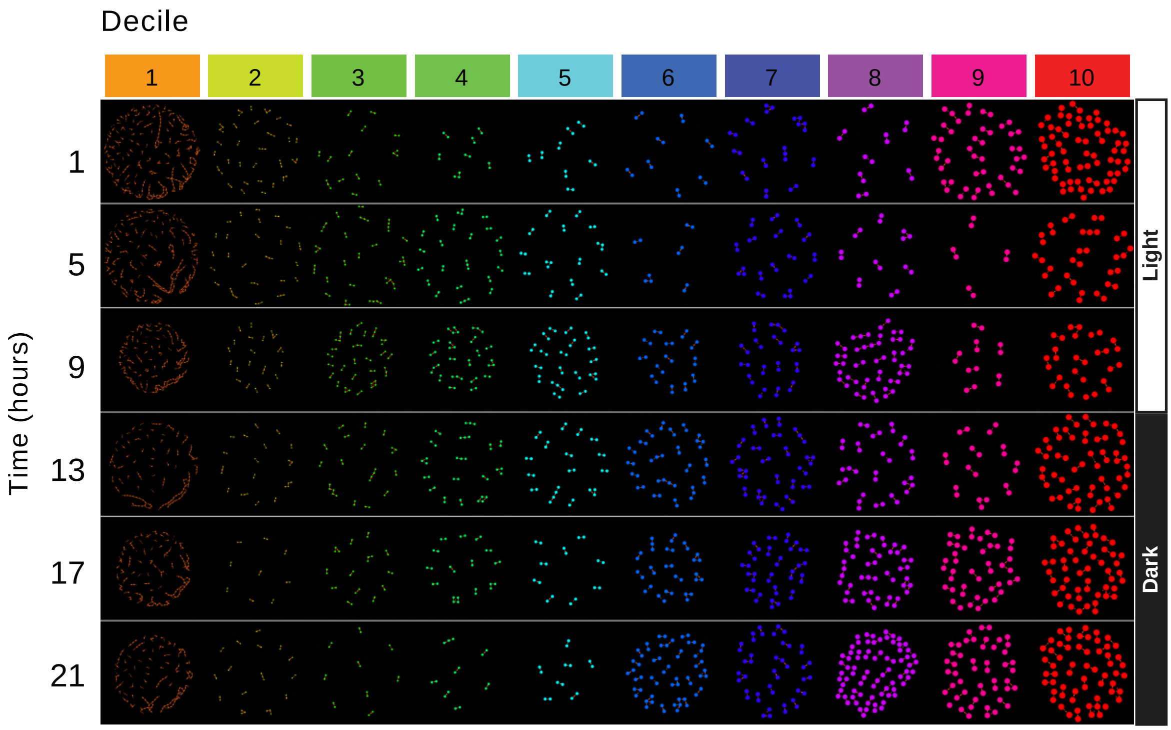

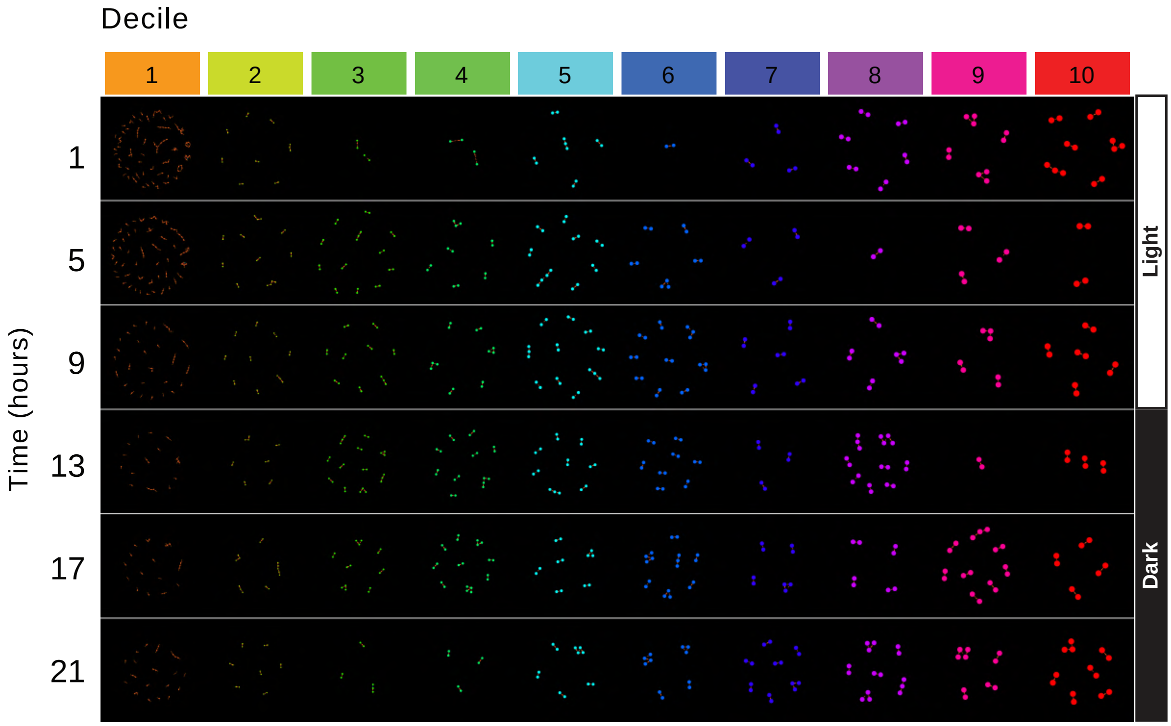

### Supplementary Figure 7

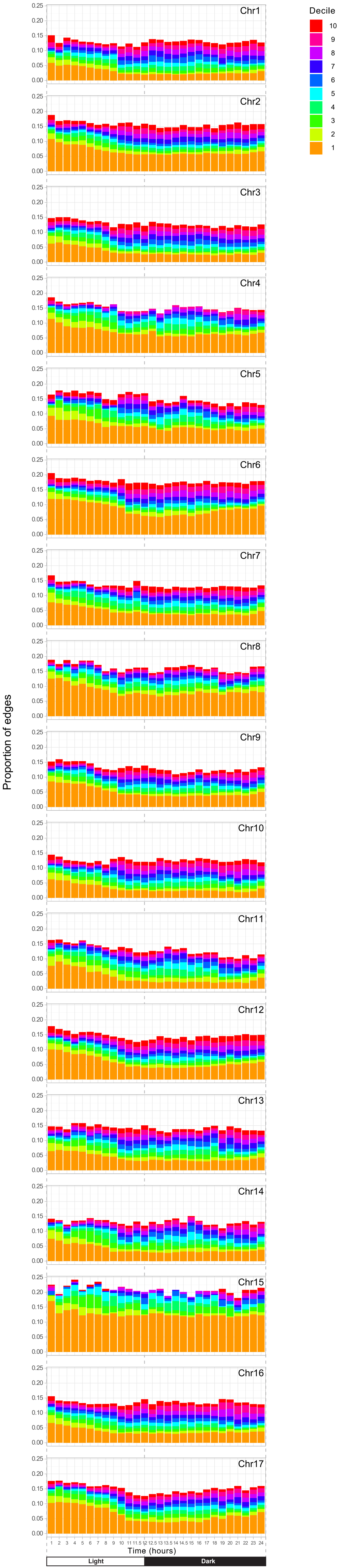

### Supplementary Figure 8

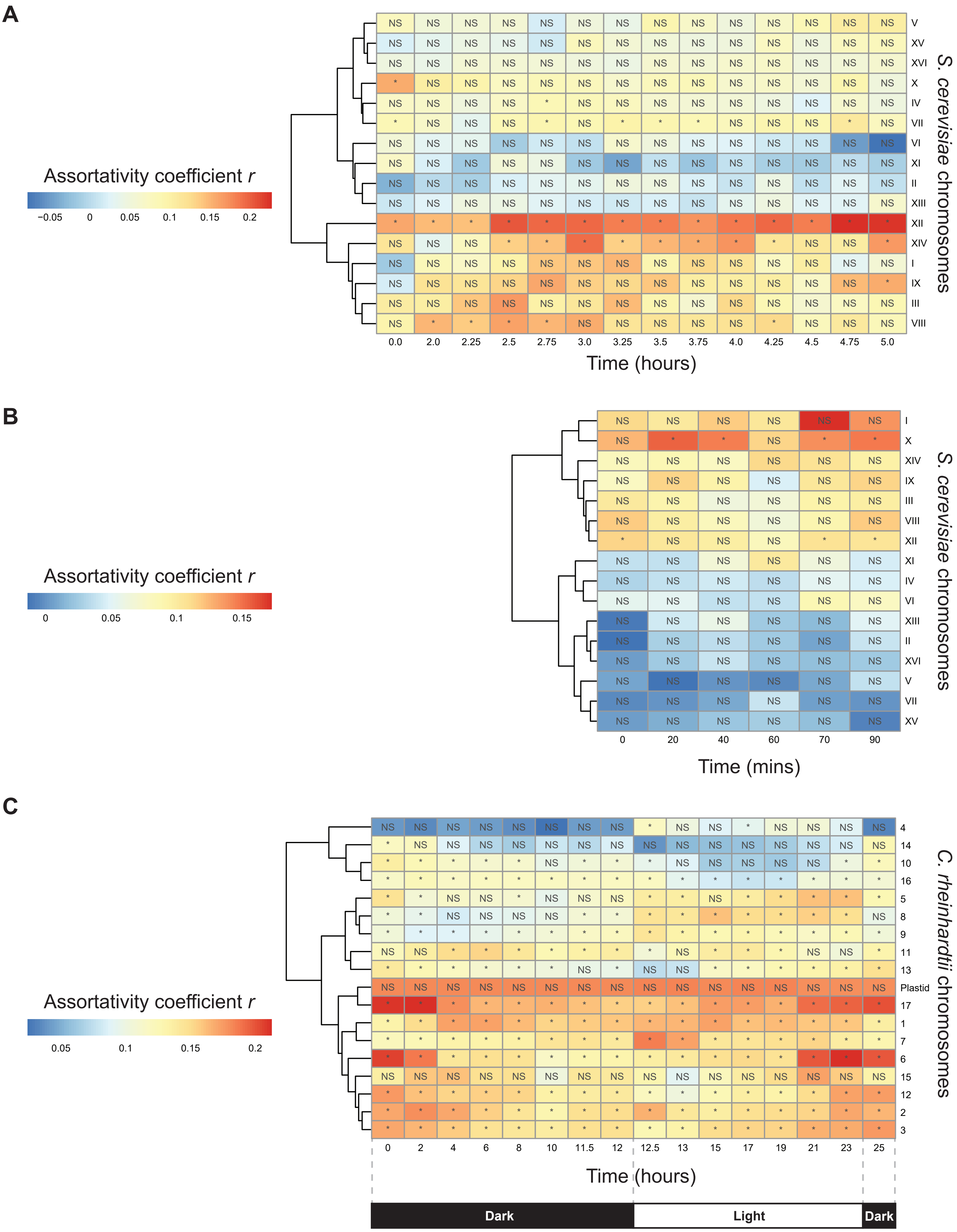
