## Supplementary Figure Legends for "GenDiMug: a generic inclusive multigraph framework to analyze genomic organizations and -omics data at multiple scales"

### Supplementary Table S1. Size and density of the networks built from 23 genomic datasets.

All datasets comprised the features annotated on the genome in the indicated assembly, as well as inferred features from these annotations (intergenic and terminal regions, introns) and additional features identified by external tools (protein domains, DNA repeats).

### Supplementary Figure S1. Metrics used to measure labelled nodes proximity in the network

(A) The assortativity coefficient *r*, as defined by Newman (14), measures the assortative mixing of nodes in a graph based on their attributes. In the formula, $e_{ii}$ is the fraction of edges in a network that connect two vertices of type *i*, $a_i$ and $b_i$ are the fractions of each type of end of an edge (in a directed graph; in undirected graphs, a = b) that is attached to vertices of type *I*, and $\Sigma_i$ means a sum for each type *I*. If the node attributes are numeric, as is the case e.g. with transcriptomic data, *r* is the Pearson correlation coefficient of the specified numeric attribute across edges. (B) Toy examples of disassortative and assortative networks for two node labels A and B; r can take values comprised between -1 (for a completely disassortative network) and 1 (for a completely assortative network), a value of 0 indicating random mixing of nodes. (C) The shortest path between two similarly labelled nodes is the minimal number of edges connecting these two nodes (length = 3 for the provided toy network).

### Supplementary Figure S2. Relationship between length of shortest paths and distance

Violin plots displaying the distribution of genomic distances associated to the shortest paths of indicated length (ranging from 1 to 50 edges) in the gene-only spatial networks of *S. cerevisiae*, *S. pombe*, *C. reinhardtii*, *C. elegans*, *D. melanogaster*, *H. sapiens*, *M. musculus*, *E. coli* and *S. islandicus*.

### Supplementary Figure S3. Shortest paths analysis in prokaryotic gene-only spatial networks labelled for gene essentiality

(A, B) Number of direct edges (shortest paths of length 1) and (C, D) average shortest path length between essential genes in (A, C) *E. coli* and (B, D) *S. islandicus*, before (light grey) and after (dark grey) correction for known sources of gene clustering. The reference values are indicated by squares and box plots represent the distributions of random values in label permutation tests. A star indicates when the reference value is significantly higher (A, B) or lower (C, D) than random values ($p < 0.05$ after Bonferroni correction, label permutation test).

### Supplementary Figure S4. Shortest paths analysis in functionally annotated prokaryotic gene-only spatial networks.

(A, B) Number of direct edges (shortest paths of length 1) and (C, D) average shortest path length between similarly annotated genes in (A, C) *E. coli* (COG functional annotation) and

(B, D) *S. islandicus* (ArCOG functional annotation), before (light grey, top plot panels) and after (dark grey, bottom plot panels) correction for known sources of gene clustering. The COGs/ArCOGs functional categories are ordered by functional classes, namely INFORMATION STORAGE AND PROCESSING (J: Translation, ribosomal structure and biogenesis. A: RNA processing and modification. K: Transcription. L: Replication, recombination and repair. B: Chromatin structure and dynamics); CELLULAR PROCESSES AND SIGNALING (D: Cell cycle control, cell division, chromosome partitioning. V: Defense mechanisms. T: Signal transduction mechanisms. M: Cell wall/membrane/envelope biogenesis. N: Cell motility. Z: Cytoskeleton. W: Extracellular structures. U: Intracellular trafficking, secretion, and vesicular transport. O: Post-translational modification, protein turnover, chaperones. X: Mobilome: prophages, transposons.); METABOLISM (C: Energy production and conversion. G: Carbohydrate transport and metabolism. E: Amino acid transport and metabolism. F: Nucleotide transport and metabolism. H: Coenzyme transport and metabolism. I: Lipid transport and metabolism. P: Inorganic ion transport and metabolism. Q: Secondary metabolites biosynthesis, transport and catabolism.). The POORLY CHARACTERIZED class (R: General function prediction only. S: Function unknown.) have been omitted from the analysis. The reference values are indicated by squares and box plots represent the distributions of random values in label permutation tests. A star indicates when the reference value is significantly higher (A, B) or lower (C, D) than random values ($p < 0.05$ after Bonferroni correction, label permutation test).

**Supplementary Figure S5. Intersection of gene co-expression and genomic networks**

(A) The gene-only network of S. islandicus with known operons represented by colored square nodes: (inset 1) NADH dehydrogenase (red) and S-layer (yellow) operons; (inset 2) ATP-synthase operon (green); (inset 3) archaellum (blue) and succinate dehydrogenase operons (pink). (B-D) Edges of gene co-expression networks, connecting nodes representing genes with significant positive co-expression (Gomez-Raya-Vilanova et al., 2025), were intersected with the spatial edges of the gene-only network of S. islandicus to obtain the sets of spatially clustered and co-expressed genes for the cell cycle (B) G1 (174 components, 486 nodes, 318 edges), (C) S (162 components, 401 nodes, 239 edges) and (D) G2 (176 components, 444 nodes, 268 edges) phases.

**Supplementary Figure S6. Clusters of iso-expressed genes in *C. reinhardtii* labelled gene-only networks**

Sorted lists of connected components (iso-expressed gene nodes connected by edges) from the gene-only networks for *C. reinhardtii* (A) chromosomes 6 (significantly assortative gene expression in the dark) and (B) 14 (weakly assortative gene expression) from sample 1 of the Zones et al, 2015 dataset, with nodes colored according to each decile of gene expression, as in Supplementary Fig S7 and Supplementary Movie S1, corresponding to transcriptomic data at time points 1, 5, 9, 13, 17 and 21 hours (44). Node size also reflects the level of expression, from decile 1 (on the left, smallest nodes) to decile 10 (on the right, largest nodes). Light and Dark conditions of the time-course are indicated on the right.

**Supplementary Figure S7. Proportions of edges linking genes in the same expression decile in *C. reinhardtii***

From a dataset produced by sampling synchronized cultured cell populations of the unicellular green alga *C. reinhardtii*, every hour or half-hour over a circadian cycle with a 12 h photoperiod, indicated with the boxes Light and Dark (44), the deciles of expression derived from normalized read count values were used as node numeric labels in the corresponding gene-only network. For each chromosome, stacked bar plots (colored according to each decile of expression, as indicated) display the changes in proportions of edges linking nodes belonging to the same decile (calculated over the total number of edges in the chromosome network) over time.

**Supplementary Figure S8. Additional assortativity analysis of gene-only spatial networks labelled for gene expression.**

(A-C) Heatmaps displaying the values of the assortativity coefficient r computed for numeric node labels corresponding to centiles of expression, computed from the distribution of normalized read counts in RPKM (A, B) or FPKM (C). The gene networks used for each chromosome and each time point were labelled using transcriptomic datasets produced by sampling synchronized cultured cell populations of (A, B) the yeast S. cerevisiae, (A) at the indicated time points during meiosis (GSE90008, unpublished > RPKM), and (B) every 20 or 10 minutes during the cell-cycle (GSE 240513, sample AF1 > RPKM) (Litsios et al., 2024), and (C) the unicellular green alga C. reinhardtii, every hour or half-hour over a circadian cycle with a 12 h photoperiod, indicated with the boxes Dark and Light (GSE112394 > FPKM) (Strenkert et al., 2019).

**Supplementary Movie S1. Dynamics of *C. reinhardtii* gene expression in labelled gene-only networks**

Visualization of parts of the gene-only networks for *C. reinhardtii* chromosomes 6 (significantly assortative gene expression in the dark) and 14 (weakly assortative gene expression) from sample 1 of the Zones et al. 2015 dataset, with nodes colored according to each decile of gene expression, as in Figs S6, S7, corresponding to transcriptomic data at time points 1, 5, 9, 13, 17 and 21 hours (44). Node size also reflects the level of expression, from decile 1 (smallest nodes) to decile 10 (largest nodes). Light and Dark conditions of the time-course are indicated on the left.
